# Determination of Capsaicin content and pungency levels in Dalle Khursani accessions, a polyploid *Capsicum* species specific to Darjeeling-Sikkim Himalaya, India

**DOI:** 10.1101/2024.09.25.614886

**Authors:** Tulsi Saran Ghimiray, Biswajit Majumdar, Sourav Man

## Abstract

The genus Capsicum includes more than 30 species, of which only five are cultivated. The pungency in chillies is primarily due to the accumulation of capsaicinoids such as capsaicin and dihydrocapsaicin, which varies significantly among species and varieties. Dalle Khursani, a unique landrace of Capsicum sp specific to the Darjeeling-Sikkim Himalaya in India, is highly valued for its pungency. However, very limited research exists on the variation of capsaicin content across different accessions of Dalle Khursani.

The present study was undertaken to evaluate the capsaicin content and pungency levels of seven accessions collected from diverse altitudinal locations. High-Performance Liquid Chromatography (HPLC) was used to quantify capsaicin content.

The concentration of capsaicin varied between 19.05 mg/g to 28.54 mg/g with the corresponding Scoville Heat Units (SHU) ranging from 304,863 to 456,636 revealing the presence of significant variations for pungency within the accessions of Dalle Khursani that may be attributed to genetic or environmental factors. Accessions DK-02 and DK-03 exhibited the highest capsaicin levels despite differing altitudinal growing environments. The results suggest a strong genetic influence on capsaicinoid content, with environmental factors playing a lesser role. For the first time, the capsaicin content and pungency levels in Dalle Khursani accessions are reported in this study.

## INTRODUCTION

Capsicum is one of the most consumed spices that have been cultivated for thousands of years. The genus *Capsicum*, belonging to the *Solanaceae* family, includes more than 30 species of which only five of them (*Capsicum annum* L., C. *chinense* Jacq., C. *frutescens* L., C. *baccatum* L. and C. *pubescens)* are cultivated.

The heat of chilli is due to the accumulation of capsaicinoids, a group of lipophilic alkaloids unique to *Capsicum* (Nelson & Dawson, 1923). The two most abundant capsaicinoids in peppers are capsaicin (8-methyl-N-vanillyl-trans-6-nonenamide) and dihydrocapsaicin (8 methyl-Nvanillylnonanamide), both constituting about 90%, with capsaicin accounting for ∼71% of the total capsaicinoids in most of the pungent varieties (Kosuge & Furuta, 1970). The Scoville Heat Test, an organoleptic evaluation introduced by Scoville in 1912, has traditionally gauged pungency levels. Over time, various methods for identifying and quantifying capsaicinoids have emerged, with High-Performance Liquid Chromatography (HPLC) considered as the most reliable and rapid technique (Collins *et al*., 1995; Peusch *et al*., 1997).

Landraces, defined as variable plant populations adapted to local agro-climatic conditions are named, selected, and maintained by traditional farmers to fulfil their social, economic, cultural, and ecological needs (Teshome *et al*., 1997). Dalle Khursani, a distinctive Chilli landrace is believed to be indigenous to Sikkim and the hill districts of Darjeeling and Kalimpong in West Bengal and reported to be a polyploid in nature (Jha *et al*., 2017), and has obtained a Geographical Indication (GI) in 2018.

No literature exists regarding the variations in capsaicin content of Dalle Khursani across different accessions in the two hill districts of West Bengal, India. The current study aims to investigate variations in capsaicin content and pungency levels among seven accessions of Dalle Khursani.

## MATERIALS AND METHODS

The entire work was carried out at the Central Instrument Centre and Quality Control Laboratory, Uttar Banga Krishi Viswavidyalaya, Pundibari, West Bengal, India. Fruits of Dalle Khursani were collected from seven different altitudinal locations at the fully ripened stage (Table.1). As this typical landrace is generally consumed fresh with seeds, the entire fruit along with seeds and placenta was considered for determining the capsaicin content.

**Table 1.**
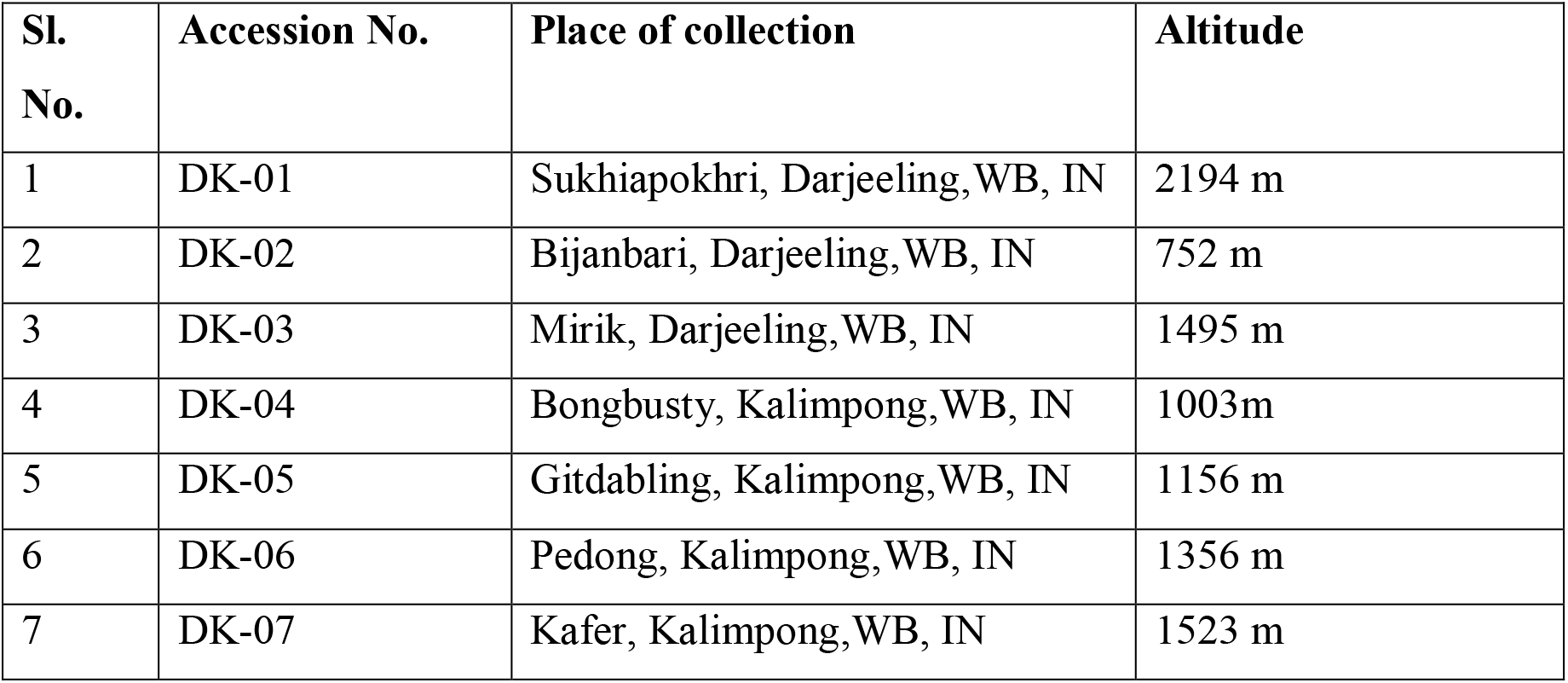
List of Dalle Khursani accessions and its place of collections.

Standard capsaicin (12084-10G) was obtained from Sigma Aldrich Co. (USA) and used as external standards. HPLC-grade methanol and acetonitrile were used. HPLC-grade water was used throughout the experiment. An HPLC system (Waters Corp., Milford, MA) with a single 515 solvent delivery pump and a 2489 UV/Vis Detector was used for high-throughput isocratic analyses. The Novapak C18 column (150 mm × 4.6 mm, 5 μm) was maintained at 30°C. The mobile phase was isocratic with 70% solvent B (100% methanol) and 30% solvent A (10% methanol solution v/v) with a flow rate of 1 mL/min for 10 minutes. The mobile phase was run through the system for 45 minutes to equilibrate the column before injecting 20μl of the sample. The UV detection wavelength was set at 280 nm as it is the maximum absorbance of capsaicinoids.

The capsaicin standard was used to obtain a calibration curve based on the peak area ratio for the known concentration of the external standard. A stock solution of the standard was prepared by dissolving 5 mg of capsaicin in 5 ml of acetonitrile and further diluted to desired concentrations in acetonitrile to generate the calibration curve.

The fruits without visible damage were selected, washed with double distilled water, and dried. Extractions and quantifications of capsaicin for all the accessions were done in triplicate using a modified HPLC procedure for the short-run method as described by Collins et al. (1995). Acetonitrile, chosen in the present investigations for its high extraction ability (Karnka *et al*., 2002), was used to extract 1g of dried chilli powder for each sample in 10 ml of solvent. The samples were placed in a water bath at 80°C for four hours, with occasional swirling at hourly intervals, and then allowed to cool at room temperature. About 2 ml of supernatant from each sample was extracted and filtered through a polyethylene membrane of 0.45 μm of pore size and stored at 5 °C in a refrigerator until analysis.

The determination of capsaicin content in the sample extracts was performed using external standards. The standard solutions were run on the HPLC and the standard curves were generated by plotting peak area against concentration. Three replicates were performed for each of the seven accessions. Capsaicin concentration was expressed in mg/g of dried weight. ANOVA and LSD test were used for the statistical analysis of the results and analysed using Grapes version 1.1.0 software (Gopinath *et al*., 2021). Capsaicin content was converted into Scoville Heat Units (SHU) based on concentration values (Helrich, 1998).

## RESULTS AND DISCUSSION

Significant variation was noted for capsaicin content among the accessions studied (Table. 2). The concentration of capsaicin varied between 19.05 mg/g to 28.54 mg/g (Fig. 1A) with corresponding SHUs from 304,863 to 456,636 (Fig. 1B) with DK-03 exhibiting the highest concentration of capsaicin closely followed by DK-02. It was interesting to note that capsaicin content in these two accessions did not vary significantly even though it differed widely in its growing environment, while differing significantly with rest of the collections. The results indicated the presence of significant variations for pungency within the accessions of Dalle Khursani, which may be attributed to genetic factor with the micro-climatic factors, soil composition and water availability contributing to pungency.

**Fig. 1.**
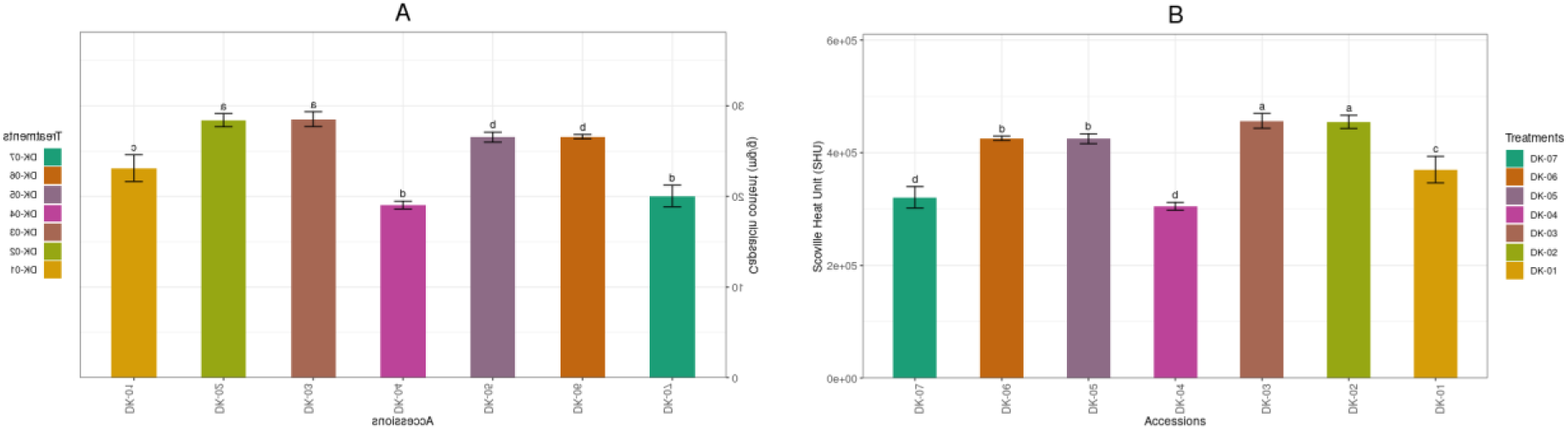
Determination of capsaicin content (A) and Scoville Unit (B) in seven accessions of Dalle Khursani.

**Table 2.** Capsaicin content and pungency level in seven accessions of Dalle Khorsani (Capsicum annuum)

| Accessions | Capsaicin (mg/g) | Scoville Heat Units (SHU) | Level of pungency |
| --- | --- | --- | --- |
| DK-01 | 23.127±0.676c | 370,015±10813.385c | Very highly pungent |
| DK-02 | 28.423±0.336a | 454,805±5367.042a | Very highly pungent |
| DK-03 | 28.540±0.376a | 456,636±6042.695a | Very highly pungent |
| DK-04 | 19.050±0.197d | 304,863±3161.966d | Very highly pungent |
| DK-05 | 26.543±0.251b | 424,741±4032.506d | Very highly pungent |
| DK-06 | 26.607±0.110b | 425,721±1778.537b | Very highly pungent |
| DK-07 | 20.053±0.546d | 320,869±8744.494b | Very highly pungent |
| Range | 19.050-28.540 | 304,863-456,636 |  |
| Mean | 24.620 | 393,950 |  |
| SEm (±) | 0.430 | 6889.957 |  |
| CD (0.05) | 1.325 | 21230.071 |  |
| CV | 3.026 | 3.029 |  |

Note: Each value is an average of three samples (±) standard error of the mean. The means followed by different letters in the same column are significantly different (p ≤ 0.05) by LSD test.

The capsaicinoid content in hot peppers is reported to be a genetically controlled trait strongly influenced by the environment and the fruit developmental stage (Burgos-Valencia et al., 2020; Hernandez-Pérez et al., 2020; Zewdie & Bosland, 2000). The pungency characteristics can be extremely variable and highly sensitive to the growing conditions of the plant and can be caused by variations in growing conditions or maturity (Cordell & Araujo, 1993; Govindarajan & Sathyanarayana, 1991; Todd et al., 1977). Iwai et al. (1979) had stated that peppers from the same plant can vary in their capsaicinoid profiles merely due to differences in postharvest ripening conditions.

The Capsaicin content and Scoville heat unit (SHU) for all seven accessions are presented in Table 2. The capsaicin contents obtained from different fruits were converted to SHU to classify them according to their various pungency levels. The capsaicin content was converted from parts per million (ppm) to SHU as per the method given by Helrich (1998). All the accessions of Dalle Khursani analysed in the present study can be classified as very highly pungent as the Scoville Heat Units (SHU) values exceed 80,000 (Weiss, 2002). A representative chromatogram depicting the distinct peak of capsaicin from a Dalle Khursani accession (DK-03 collected from Mirik, Darjeeling) is presented in Fig. 2.

**Fig. 2.**
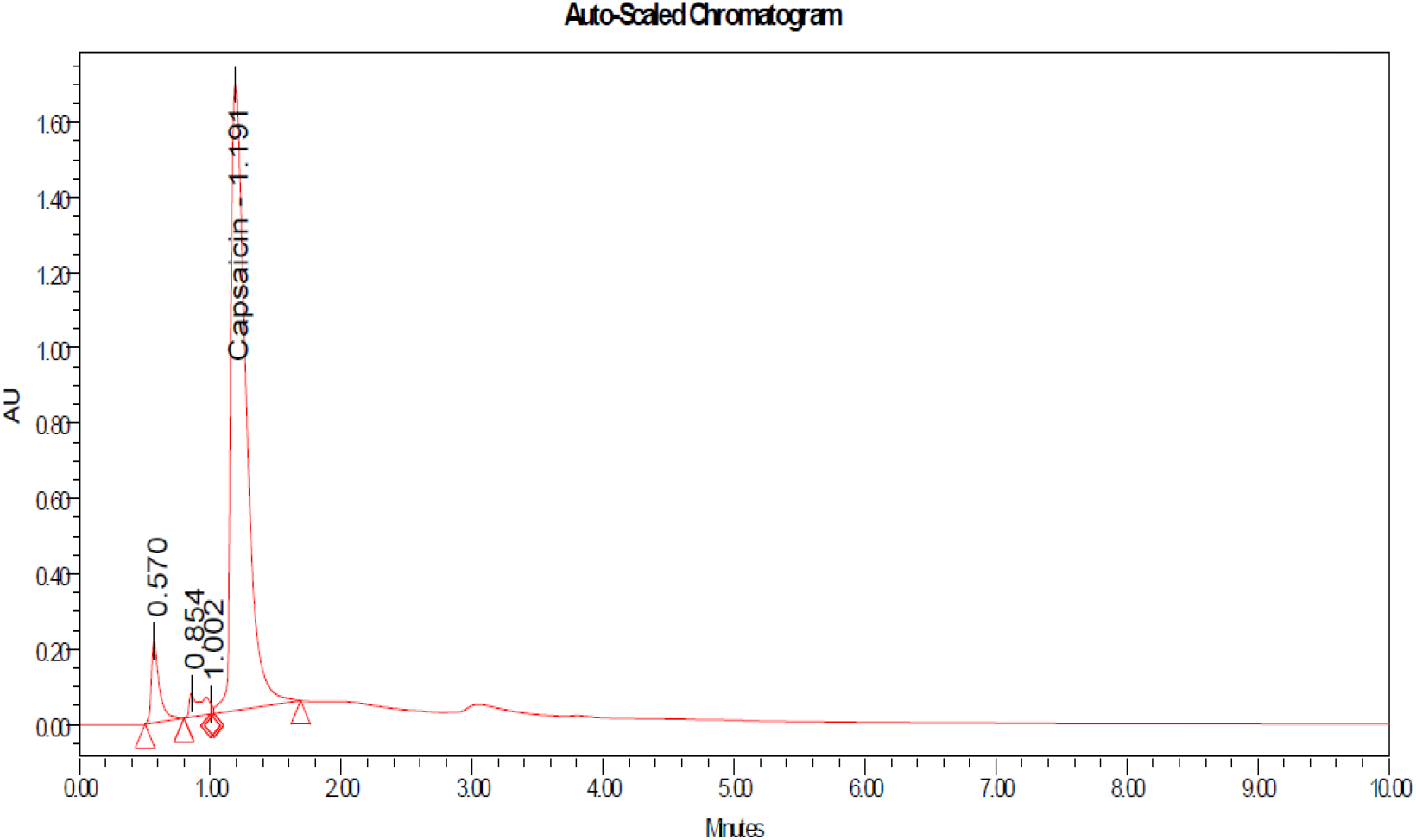
Chromatogram of capsaicin content in Dalle Khursani accession (DK-03)

Available literature suggests that *Capsicum chinense* species includes the most pungent chilli peppers (Bosland & Baral, 2007; Canto-Flick *et al*., 2008) and some authors have also mentioned Dalle Khursani as *Capsicum chinense*. But recent works by Jha et al. (2017) indicated its closer affinity to *Capsicum annuum*, and have reported the polyploid nature of this landrace. Since this landrace is specific to Darjeeling-Sikkim Himalaya, all the accessions showed high level of pungency and selection may be imposed upon these lines based on the yield and other biochemcial parameter. There is also a need for genetic improvement of this typical landrace for the development of a stable variety.

Available literature on the pungency level of Dalle Khursani are scanty and reported to be in the range of 100,000 to 350,000 SHU (Lepcha *et al*., 2023) similar to that of Habanero pepper. In contrast, we have observed considerable variation in pungency levels for the seven accessions evaluated, much higher than those already reported. The capsaicin content and corresponding pungency level of these accessions can be expected to have a higher level if only the placenta have been used for the analysis as capasaicin is produced in placental tissues.

## CONCLUSION

Considerable variation in capsaicin content was observed among the seven accessions of Dalle Khursani. However, no clear relationship between the growing environment and capsaicin content was found. The present study reports for the first time the capsaicin content and variation in pungency level in Dalle Khursani over different altitudinal locations in the two hill districts of West Bengal, India. It is interesting to note that being a polyploid and being grown in a specific region in India, further studies need be emphasised on genetic improvement of this landrace since they are prone to various pest and disease that cause considerable loss to the growers.

